# Minimal circuit motifs for second-order conditioning in the insect mushroom body

**DOI:** 10.1101/2023.09.11.557174

**Authors:** Anna-Maria Jürgensen, Felix Johannes Schmitt, Martin Paul Nawrot

## Abstract

In well-established first-order conditioning experiments, the concurrence of a sensory cue with reinforcement forms an association, allowing the cue to predict future reinforcement. Once a sensory cue is established as a predictor, it can also serve as indirect reinforcement, a phenomenon referred to as second-order conditioning. In the insect mushroom body, such associations are encoded in the plasticity of the synapses between the intrinsic and output neurons of the mushroom body, a process mediated by the activity of dopaminergic neurons that encode reinforcement signals. In second-order conditioning, a new sensory cue is paired with an already established one that presumably activates dopaminergic neurons due to its predictive power of the reinforcement. We explore minimal circuit motifs in the mushroom body for their ability to support second-order conditioning. We found that dopaminergic neurons can either be activated directly by the mushroom body’s intrinsic neurons or via feedback from the output neurons via several pathways. We demonstrate that the circuit motifs differ in their computational efficiency and robustness and suggest a particular motif that relies on feedforward input of the mushroom body intrinsic neurons to dopaminergic neurons as a promising additional candidate for experimental evaluation. It differentiates well between trained and novel stimuli, demonstrating robust performance across a range of model parameters.

## Introduction

By forming associations between sensory cues and reinforcement during classical conditioning (first-order conditioning, FOC), animals can learn to predict the emergence of environmental factors relevant to their survival. Once a sensory cue has been established as a predictor of such reinforcement, it can act as reinforcement in second-order conditioning (SOC). SOC has been observed across species with experiments conducted in *Drosophila* [1–4] and other invertebrate [5–8] as well as vertebrate [9–11] species. SOC experiments involve two initially neutral stimuli (stimulus 1 and stimulus 2). Stimulus 1 is first paired directly with reinforcement during FOC, whereby it acquires a valence as a cue for reinforcement. Afterward, stimulus 2 is paired with stimulus 1 (SOC), causing an expansion of the acquired valence of stimulus 1 onto stimulus 2, without stimulus 2 itself being paired with the reinforcer. Afterward, both stimuli initiate a behavioral response based on their acquired valence.

In *Drosophila* [12–15] and other insects [15–18], the mushroom body (MB) is a crucial brain structure for learning and encoding relationships between sensory cues and reinforcement. The Kenyon cells (KC) are the intrinsic neurons of the MB, encoding the identity of sensory inputs in *Drosophila* [19–21] as well in other insects [22–25]. In both *Drosophila* [26, 27] and other insects [24, 25, 28, 29], they relay their output onto a much smaller number of MB output neurons (MBONs). Plasticity at the KC>MBON synapses allows MBONs to encode the valence of a sensory cue, according to extensive experimental evidence from *Drosophila* [30–35]. Neuromodulators, such as dopamine mediate this plasticity [36–42]. In *Drosophila*, it has been shown that either an inherently punishing or rewarding stimulus (electric shock, sugar) [1, 3, 4] or direct optogenetic activation of dopaminergic neurons (DANs) [3] can be utilized to deliver a reinforcement signal during FOC phase of such experiments to establish second-order memory later. Experiments in *Drosophila* have suggested that stimulus 1 itself causes activation of DANs or enhances it after being paired with reinforcement [3, 4, 43, 44]. This aligns with the analogous finding that DANs in the ventral tegmental of vertebrates respond to learned cues that predict upcoming reward [45]. The mechanism inducing synaptic plasticity during both FOC and SOC likely relies on DAN activation. During FOC, DANs are activated directly by the reinforcer (Figure 1 A). Following this conditioning procedure, stimulus 1 seems to indirectly activate the DANs during SOC (Figure 1 C, [3, 4, 44]). The strength of the behaviorally expressed stimulus 1 and stimulus 2 valence after SOC can be similarly strong [1, 3] or weaker [3, 4] for stimulus 2.

**Figure 1.**
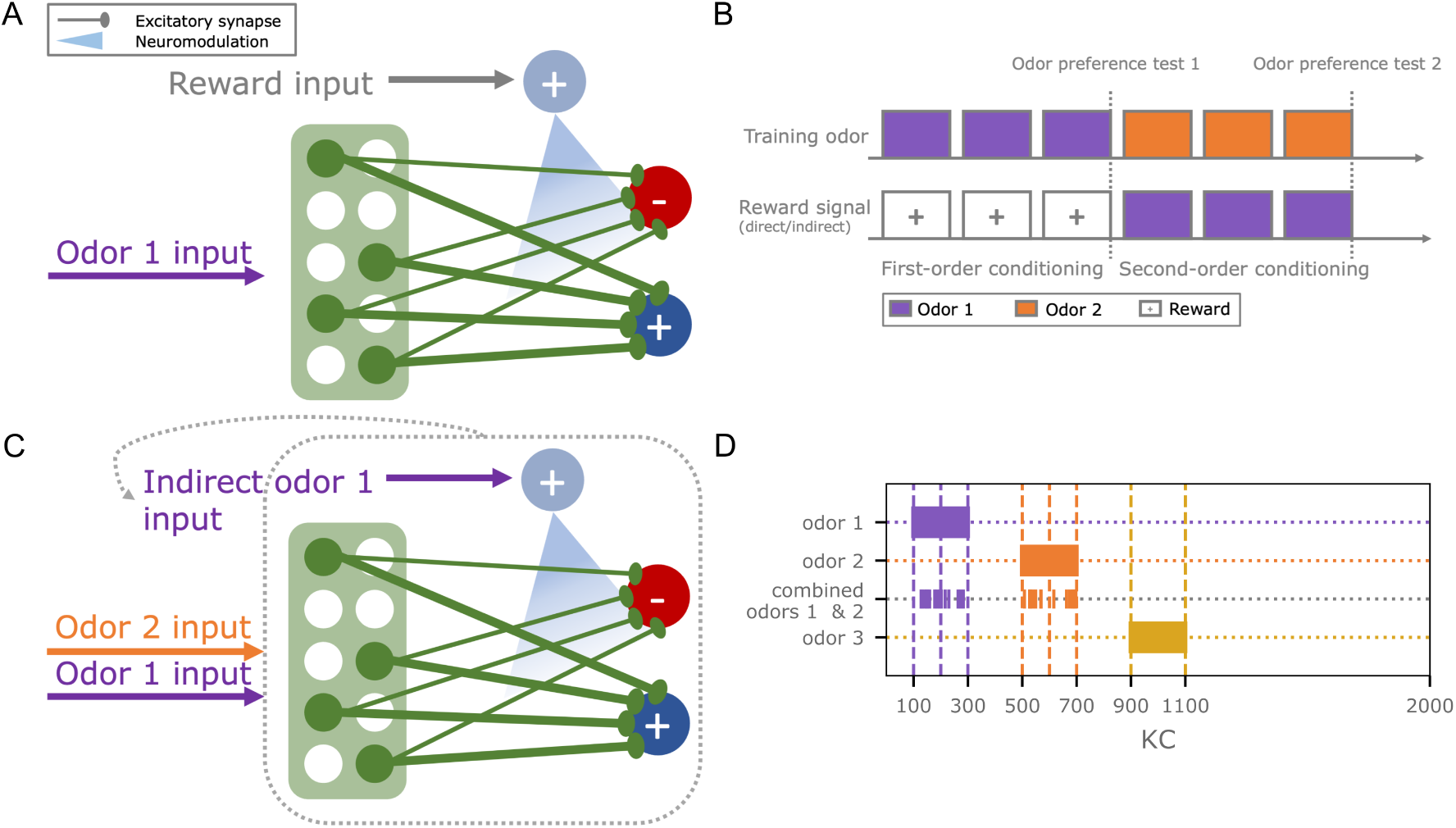
Experimental design for testing second-order conditioning. (A) Basic circuit motif for first-order conditioning, consisting of 2000 Kenyon cells (green), two output neurons (dark blue, red), and a single dopaminergic neuron (light blue). The co-occurrence of odor and reward input elicits plasticity at the MB output synapses. (B) The experimental paradigm consists of two phases (first and second-order conditioning). During first-order conditioning, odor 1 is paired with a reward. Subsequently, a novel odor 2 is paired with odor 1 during second-order conditioning. Odor valences are tested after first and second-order conditioning. (C) During second-order conditioning, the dopaminergic neuron (light blue) is indirectly activated by the previously trained odor 1 and paired with odor 2. We test different candidate mechanisms for this indirect activation of the dopaminergic neuron via the Kenyon cells (green) or the mushroom body output neurons (dark blue, red). (D) Initially, three non-overlapping odors were used in the experiments. Odors are encoded as Kenyon cell activity patterns. The joint presentation of odor 1 and odor 2 during second-order conditioning retains a randomly chosen 50% of the individual odor representations to maintain the same overall activation as with individually presented odors.

Two different circuit mechanisms lend themselves to achieving such post-FOC activation of DANs by stimulus 1: Firstly, a stimulus 1 representation among the KCs could serve as direct stimulus-specific input to the DANs [46–51]. Alternatively, the input could be supplied via MBON feedback [46, 47, 51, 52]. Their response to stimulus 1, altered by learning, could serve as a manifestation of stimulus-specific valence.

Here, we tested possible circuit motifs that could underly SOC in the insect MB using abstract and simplified network models inspired by the *Drosophila* olfactory pathway and the MB. Starting from a basic model of the MB, we explored different circuit configurations and their capacity to produce SOC in an olfactory learning protocol to identify promising candidates for experimental testing. To define our solution space, we assumed that learning in the MB depends on KC>MBON plasticity, mediated by a dopamine signal during FOC and SOC. Model circuits should be able to produce both FOC and SOC without generalizing associations with reinforcement unspecifically to novel stimuli. We tested all models in classical conditioning experiments and demonstrated their ability to support FOC and SOC. Additionally, we evaluated differences in their biological plausibility by quantifying robustness and discussing functional and anatomical evidence for the respective circuits. We found that a particular circuit that achieves DAN activation through excitatory KC input during SOC outperforms the other candidates and appears compatible with the MB anatomy. We suggest this circuit motif that differs from previously reported mechanisms [2–4] as an additional candidate for experimental tests.

## Results

### First-order conditioning in a basic mushroom body circuit

All models are based on rate units, each representing the activation of a single neuron. The basic network consists of 2000 KCs, each innervating two MBONs (Figure 1 A). It has been shown that MBONs receive inputs from many of the KCs in adult *Drosophila* [26, 51]. For simplicity, we started by modeling a complete KC>MBON connectivity. Some MBONs can be categorized as approach or avoidance signaling [30, 33, 34, 53]. In the model, this corresponds to MBON^+^ (Eqn. 2) and MBON-(Eqn. 3), respectively. Other types of MBONs were disregarded here. Initially, all synaptic KC>MBON weights were set to the same value (table 2). A single DAN is included in the model, which can be activated by the external input, representing a reinforcer in the environment (Figure 1 A). KC>MBON-synapses undergo plasticity whenever trial-based KC activation, driven by odor input, and DAN activation coincide. We employ a two-factor learning rule (Eqn. 4) at the KC>MBON-synapses, leading to a decrease in the synaptic weights with the limitation that they can not take on a negative value. DAN activation (Eqn. 1) is the sum of the model-specific external input rate *Ip*_*ext*_ (table 3) that represents reinforcement and the network internal input *Ip*_*int*_, provided via the different circuit mechanisms. In all equations, 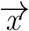 always denotes a vector. *R* represents the activity of a neuron, which can be interpreted as a spike rate of a neuron or a vector of neurons in the case of KCs, and *w* denotes a synaptic weight or a vector of weights. *LR* refers to learning rates.

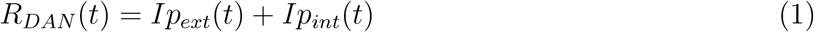

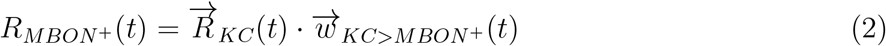

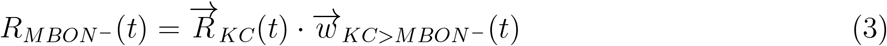

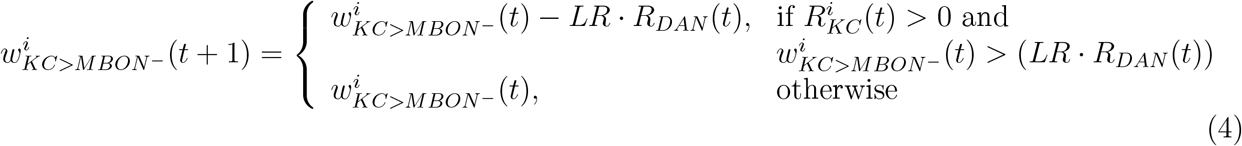

### Candidate circuits for second-order conditioning

Using the basic circuit model (Figure 1 A) as a starting point, we implemented five different extended versions of it (Figure 2). These models either rely on some form of KC>DAN input (model 1, model 2) or MBON>DAN feedback (model 3, model 4, model 5) as a means to expand the learned association of odor 1 with reinforcement to odor 2 during SOC. Unless specified otherwise, the DAN is not spontaneously active. The equations for all models can be found in table 1.

**Table 1:**
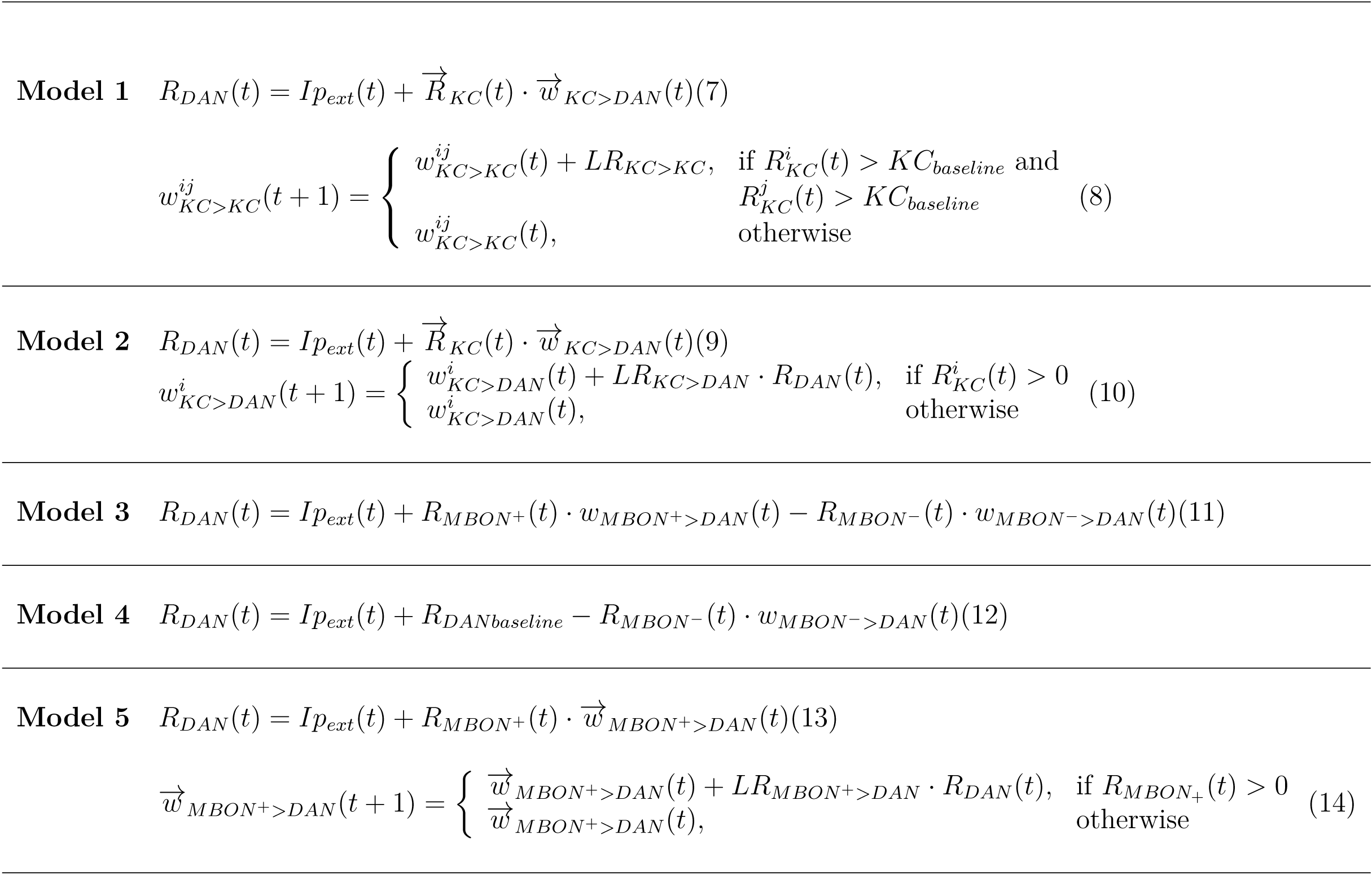
Equations underlying the different models.

**Figure 2.**
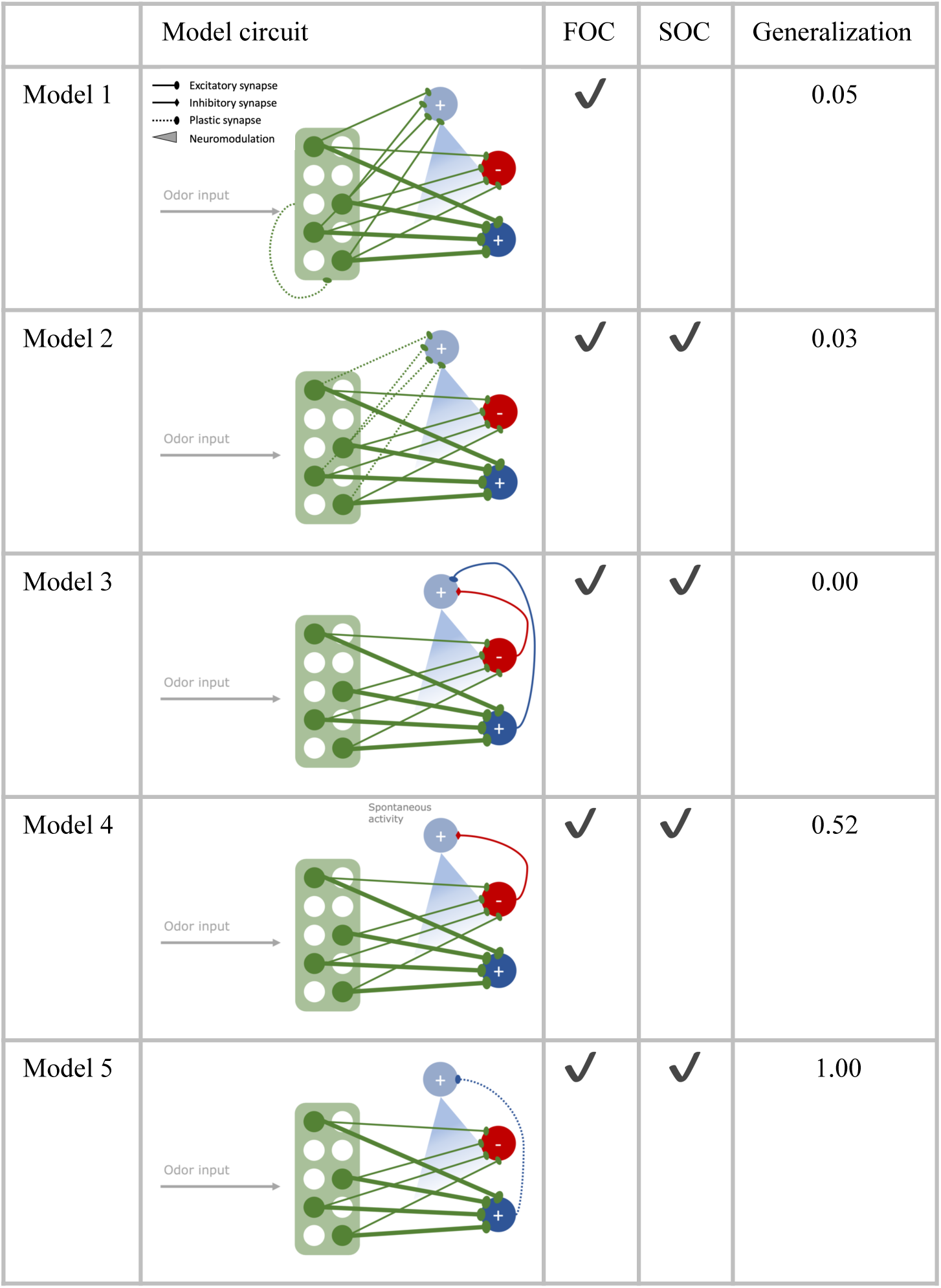
Second-order conditioning in different circuit motifs. Five different circuits were tested for their performance in first (FOC) and second-order conditioning (SOC) and the extent to which the odor-reward association generalizes to another novel odor. All circuits are constructed with 2000 Kenyon cells (green), two mushroom body output neurons (dark blue, red), and a single dopaminergic neuron (light blue), targeting the synapses between Kenyon cells and mushroom body output neurons. Additional feed-forward connections from the Kenyon cells (model 1, model 2) or feedback connections from the mushroom body output neurons onto the dopaminergic neuron (model 3, model 4, model 5) are implemented in the different circuits.

To compare the different circuit motifs in an unbiased manner, their parameters were optimized using grid search (see Methods), except the fixed parameters shared between all models (Table 2), which were kept constant to allow better comparison between the candidate mechanisms for SOC implemented in the different circuits. The goal for parameter optimization was to identify parameter combinations for each model that yield the best learning performance in an associative learning experiment that consisted of a combination of FOC and SOC learning trials (Figure 1 B). In insect learning experiments, forming a direct or indirect association with reward leads to approach behavior that can manifest in the movement toward the source of an odor or feeding-related behavior [3, 5, 6] or the avoidance of a punishment-associated odor [4]. In our model experiments, the successful acquisition of an association with reward was quantified as the normalized difference between MBON output rates (*R*_*MBON*+_ and *R*_*MBON−*_), which we will refer to as the approach bias (Eqn. 5) because MBON activity has been shown to initiate approach or avoidance behavior [30, 33, 54].

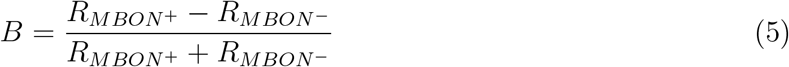

**Table 2:**
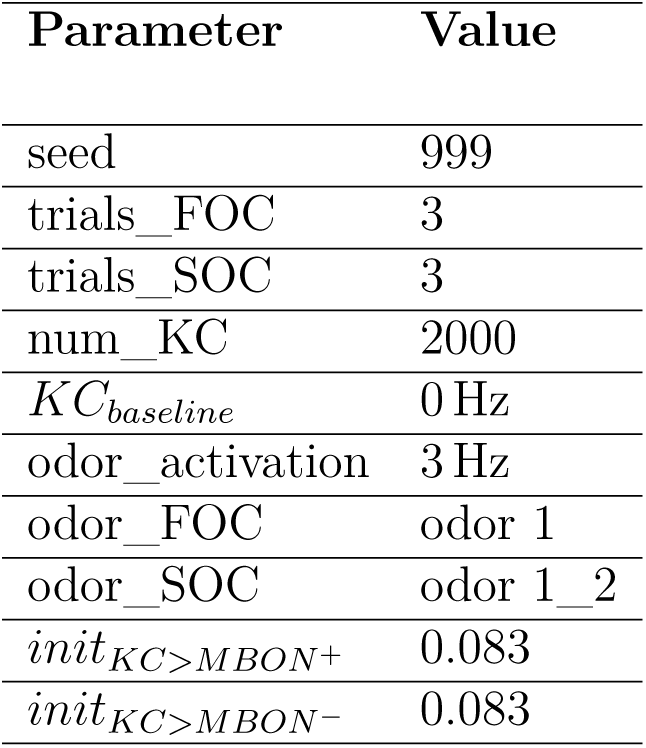
Fixed parameters shared between models.

| Parameter | Value |
| --- | --- |
| seed | 999 |
| trials_FOC | 3 |
| trials_SOC | 3 |
| num_KC | 2000 |
| $KC_{baseline}$ | 0 Hz |
| odor_activation | 3 Hz |
| odor_FOC | odor 1 |
| odor_SOC | odor 1_2 |
| $init_{KC>MBON+}$ | 0.083 |
| $init_{KC>MBON-}$ | 0.083 |

#### Model 1

Model 1 includes plastic KC>KC excitatory feedback with increments of *LR*_*KC>KC*_(table 3)), triggered by pre and postsynaptic KC activation (Eqn. 8) and KC>DAN synapses of a fixed strength (Eqn. 7).

**Table 3:** Optimized model parameters. For all models, 100 equally distributed values per parameter between the minimum and maximum values were used to construct a regular grid of parameter combinations.

| Parameter | min | max | optimum |
| --- | --- | --- | --- |
| **Model 1** |  |  |  |
| $init_{KC>KC}$ | 0 | 0 | 0 |
| $init_{KC>DAN}$ | 0.0 | 0.001 | 0.001 |
| $LR_{KC>MBON^-}$ | 0.001 | 0.004 | 0.004 |
| $LR_{KC>KC}$ | 0.0 | 0.004 | 0.000162 |
| DAN_activation | 1.0 | 10.0 | 7.272727 |
| **Model 2** |  |  |  |
| $init_{KC>DAN}$ | 0.0 | 0.001 | 0.000505 |
| $LR_{KC>MBON^-}$ | 0.001 | 0.004 | 0.00333 |
| $LR_{KC>DAN}$ | 0.0 | 0.001 | 0.000677 |
| DAN_activation | 1.0 | 10.0 | 5.727273 |
| **Model 3** |  |  |  |
| $init_{MBON^+>DAN}$ | 0.0 | 1.0 | 0.272727 |
| $init_{MBON^->DAN}$ | 0.0 | 1.0 | 0.262626 |
| $LR_{KC>MBON^-}$ | 0.001 | 0.004 | 0.003121 |
| DAN_activation | 1.0 | 10.0 | 5.00 |
| **Model 4** |  |  |  |
| $init_{MBON^->DAN}$ | 0.0 | 0.5 | 0.131313 |
| $LR_{KC>MBON^-}$ | 0.001 | 0.004 | 0.003182 |
| $R_{DANbaseline}$ | 0.0 | 30.0 | 11.212121 |
| DAN_activation | 1.0 | 10.0 | 3.727273 |
| **Model 5** |  |  |  |
| $init_{MBON^+>DAN}$ | 0.0 | 0.4 | 0.08008 |
| $LR_{KC>MBON^-}$ | 0.001 | 0.004 | 0.003182 |
| $LR_{MBON^+>DAN}$ | 0.001 | 0.01 | 0.003 |
| DAN_activation | 1.0 | 10.0 | 4.727273 |

#### Model 2

Model 2 expands the basic circuit with excitatory plastic synapses between KCs and the DAN (Eqn. 10). They are each initialized with *init*_*KC>DAN*_ (table 3) and are increased by *LR*_*KC>DAN*_ (table 3) when activation of the respective KC coincides with DAN activity, yielding DAN activation (Eqn. 9).

#### Model 3

In model 3, network input into the DAN is implemented via feedback from both MBONs (Eqn. 11). Inhibitory input with a fixed synaptic strength comes from MBON-, while excitatory input is provided by the MBON^+^ (table 3).

#### Model 4

Model 4 uses a spontaneously active DAN (*R*_*DANbaseline*_, (table 3)) in combination with inhibitory MBON->DAN input (Eqn. 12). Both effects regulate the DAN activation in the absence of reward.

#### Model 5

In model 5, an excitatory plastic MBON^+^>DAN synapse (Eqn. 14) is added to the basic circuit. When MBON^+^ and DAN activity coincide, the synaptic strength is increased by 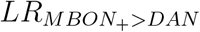 for each DAN spike (table 3). During FOC, this synapse is strengthened, allowing for activation of the DAN during SOC. This allows KCs to activate the DAN (Eqn. 13).

### Identifying optimal parameters for each model

All models were trained in a combined FOC and SOC protocol, where odors were used as stimuli and tested for their approach bias first after completing FOC with odor 1 and reward, then after completing SOC with odors 1 and 2 (Figure 1 B, Eqn. 5). Plasticity was disabled during testing to isolate odor valence acquired during the respective training trials without the influence of the test itself. Additionally, a novel odor 3 was included in both tests to examine any generalization of the reward association that might have occurred during the FOC or SOC training processes. To assess the ability of the parameters of the different circuit motifs to support SOC, we optimized each model for the highest possible SOC performance, which translates to maximizing the approach bias for odor 2 after three SOC trials. Additionally, we introduce several criteria the models must fulfill to ensure that the learning effect for odor 2 is odor-specific and originates from the respective mechanism applied during the SOC trials. These criteria are:

- post FOC odor 1 approach bias = 1
- post FOC odor 2 approach bias = 0
- post FOC odor 3 approach bias = 0
- post SOC odor 1 approach bias = 1
- post SOC odor 3 approach bias = 0

Additionally, DAN and MBON rates should never exceed 20 Hz and 50 Hz, respectively, to stay within the biologically observed range for MBONs [34] and DANs [55]. The reported DAN spike rates are responses to an electric shock that were reported to range up to 10 Hz during a 200ms stimulation. Since the response of the DANs showed a temporal delay, we decided to use 20 Hz instead.

All parameter combinations that fulfilled the above criteria for each model were further evaluated. Among all of these parameter combinations, we selected those for each model that yielded the highest SOC performance, measured by the odor 2 approach bias after SOC. We refer to each of these as an *optimal learner*. Grid search for all models 1-5 yielded several *optimal learners*. We computed the average *optimal learner* by averaging all *optimal learners* within every parameter. We argue that this average *optimal learner* approximates the center of all equally good parameter combinations. Next, Euclidean distances were computed between all z-standardized *optimal learners* and the average *optimal learner*. The parameter combination with the smallest distance to the average parameter combination in an n-dimensional z-standardized space (n=number of optimized parameters) was selected (see Methods). We assume that parameter combinations closer to the average can be considered biologically more plausible because their central location makes them more robust to parameter deviations in all directions (see Discussion).

The basic learning model (Figure 1 A) fulfilled the criteria for FOC learning, but no parameter combination could accommodate SOC, yielding no approach bias for odor 2 after SOC. There was at least one optimal parameter combination that fulfilled the optimization criteria for each extended candidate circuit (Figure 2, Figure 3 A). All models acquired an optimal approach bias of 1 for odor 1 at both test times, indicating that the association of odor 1 and reward is learned during FOC and fully retained throughout the SOC protocol. Tests with odor 3 always yielded an approach bias of 0 for all models, indicating that the approach bias does not generalize inadmissibly to fully disjunct odors. All models, except model 1, achieved equally good SOC performance, as indicated by an approach bias of 0.33 for odor 2 after SOC. Model 1 only acquired a bias of 0.02. For each model, the maximum value of SOC performance is determined by the implementation of the compound presentation of odors 1 and 2 (see Methods). The approach bias of 0.33 for SOC (Eqn. 5) is the highest value any model can achieve in this experiment. In the experimental design, 50% of KCs were activated by odor 2 during SOC. Thus, only 50% of the KC>MBON-synapses could be altered during SOC. The value of 0.33, therefore, represents the best possible SOC performance. An additional experiment was conducted as a control where the KC activation patterns for odors 1 and 2 were added during SOC (yielding 400 active KCs). The maximal performance achieved by models 2-5 was 1 in this case.

**Figure 3.**
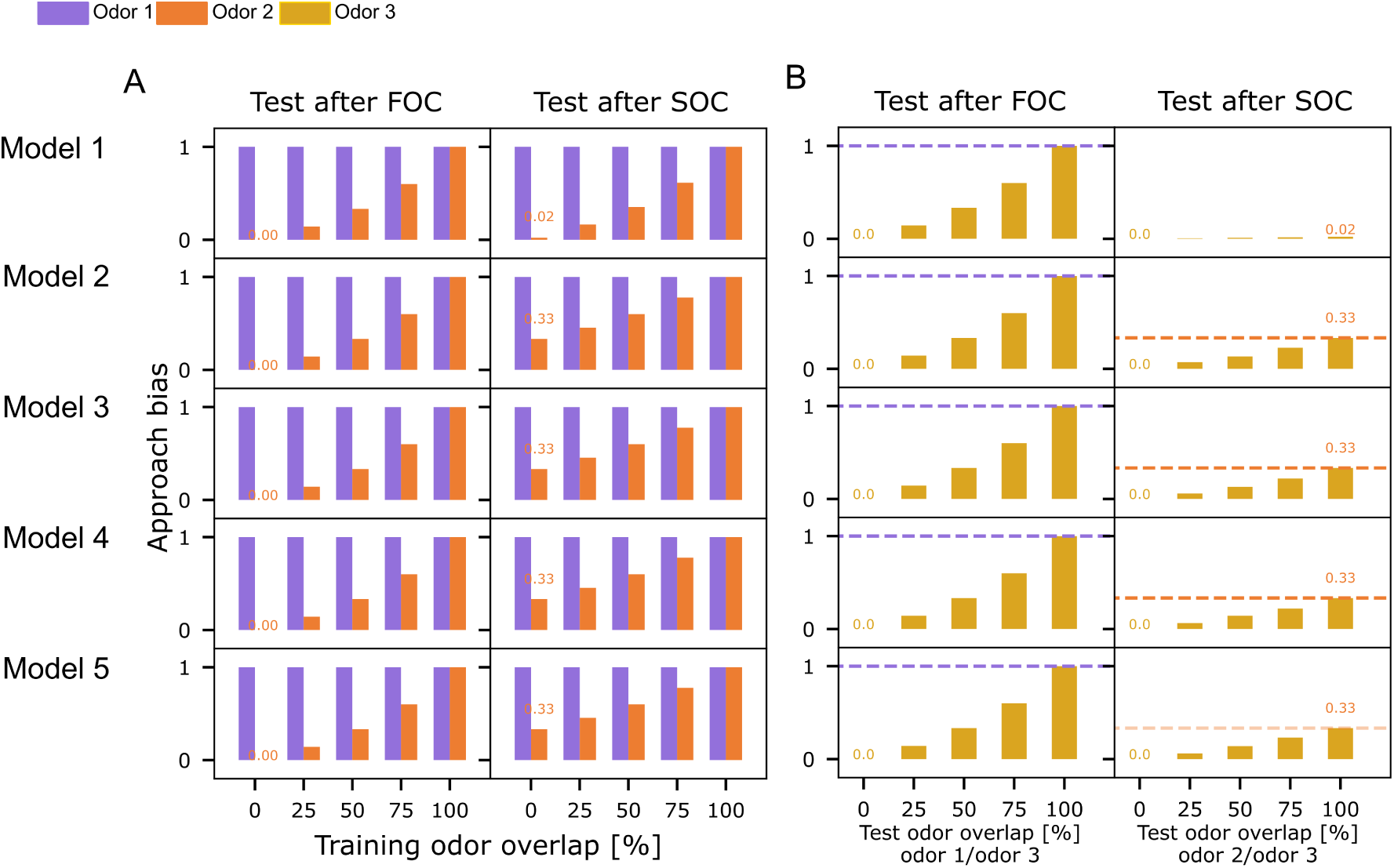
Reward generalization for overlapping training or test odors. (A) All five models were tested for their odor approach bias (Eqn. 5) to odors 1 (purple) and 2 (orange) after first (FOC) and second-order conditioning (SOC). The overlap between odors 1 and 2 was varied. (B) All models were tested for their approach bias (Eqn. 5) to odor 3, after training with non-overlapping odors 1 and 2. Overlap between odors 3 and 1 or 2 was varied, respectively. Orange and purple dashed lines indicate each model’s FOC and SOC performance from an experiment without odor overlap as a reference (always the first bar).

In none of the models any approach bias for the disjunct, novel odor 3 was observed. Additionally, we extended the experimental protocol (Figure 1 B) with three trials of presenting odor 3 alone and without any reward after SOC and included another test. Depending on the degree of specificity with which the different model circuits activate the DAN, unwanted generalization of the valence to odor 3 was observed. Models 1,2,3 outperform models 4 and 5 here (Figure 2).

### Learning generalizes with increasing odor overlap

The overlap of odors 1 and 2 during training and training was varied separately (0%, 25%, 50%, 75%, 100% overlap), encoded in the percentage of KCs jointly activated between the individually presented odor 1 and odor 2 (see Methods). Across all models, both FOC and SOC approach biases (Eqn. 5) increase with the overlap between odor 1 and odor 2 (Figure 3 A). Between highly overlapping odors, the reward association generalizes. This results in an approach bias for odor 2 after FOC, even though odor 2 was not presented during FOC (Figure 3 A). A joint presentation of odor 1 and odor 2 during SOC then leads to an even higher approach bias for odor 2 (Figure 3 A). All models acquire similar biases (Eqn. 5) for odor 2, depending on the degree of overlap, except model 1, where the SOC learning effect was always lower (Figure 3 A).

Keeping odor 1 and odor 2 fully disjunct, we varied the degree of overlap between odor 3 and either odor 1 (Figure 3 B, left) or odor 2 (Figure 3 B, right) during both tests. For all models, the approach bias for odor 3 after FOC scaled with the overlap and reached the same value as odor 1 if fully overlapping (f Figure 3 B, dashed purple lines). Testing again after SOC yielded the same results, thus not depicted here. When the overlap between odor 3 and odor 2 varied at both test times, no approach bias was observed after FOC since odor 2 is not presented during the FOC trials (results not shown). In a test after SOC, all models perform similarly concerning the upper bound of the approach bias at the magnitude reached by odor 2 Figure 3 B, dashed orange lines).

### Robustness of second-order conditioning varies across the different model circuits

In the conditioning experiments reported thus far, all model circuits, except model 1, perform equally well (Figure 2, Figure 3). To further differentiate between them, we next examined the robustness of the model’s performance to variations of their parameters using three different methods.

We first quantified the percentage of optimal parameter combinations within the searched parameter grid for each model. A real brain would likely not require a single, extremely precise combination of physiological parameters to perform any computational task, such as SOC. Since the parameters of our computational models are ultimately representations of neuronal or synaptic characteristics, the stability of SOC performance across parameter combinations could hint at the degree to which a circuit motif is biologically plausible. For each respective model, four parameters were optimized, yielding a grid with 100^4^ parameter combinations. In the case of model 1, 5.9^−5^% of parameter combinations were equally optimal. Model 2 yielded 6.81%, model 3 only 0.37%, model 4 2.11% and model 5 4.29% *optimal learner* parameter combinations. From this perspective, models 2, 4, and 5 thus seem more robust compared to models 1 and 3.

As an additional measure to assessing the optimal portion of the entire solution space of possible parameter combinations, we used a method for individually sampling the four-dimensional Euclidean parameter space for each model. The four-dimensional space for each model was standardized using the range of the grid (maximum-minimum parameter value, table 3). A four-dimensional hypersphere was positioned as the point representing the average *optimal learner*, with radius=0. We then incrementally increased the radius of the hypersphere from 0 to 1 in the standardized space with 100 linearly spaced steps and, for each increase, sampled 700 data points from its surface (see Methods, Hypersphere sampling). These sampled data points do not necessarily correspond to data points from the set of equally optimal parameter combinations found in the grid search for the respective model due to the step size used in the parameter optimization. The sampled points were transformed back into their original space and then used to simulate the respective model to examine if this parameter combination would yield FOC and SOC performance that fit the criteria for the *optimal learner*. For each radius increment, we calculated the percentage of sampled points from the hypersphere surface that exhibit the same performance as the parameter combination used as the central point. Since the parameter space was standardized for each model and the radii used were the same, the results can easily be compared between the five models. We find that the models differ in their robustness to deviations of the parameters from their optimal values. Model 2 strongly outperforms the other models. While models 4 and 5 are more robust than model 3, model 1 demonstrates no robustness. Our grid search for model 1 yielded no variability in three of the four optimized parameters (*init*_*KC>KC*_, *init*_*KC>DAN*_, *LRKC>MBON _, LR*_*KC>KC*_, Table 3).

A third approach to comparing the robustness between the different model circuits is to quantify how well they retain their FOC and SOC performance when variability is introduced into the connectivity matrix or the strength of the KC>MBON synapses. We varied either the number of input KCs into each MBON (Figure 5 A,B) or the strength of the synaptic connections while retaining full connectivity (Figure 5 C,D).

We varied the number of KCs providing input to each MBON between 25% and 100% for each model instance (*N* = 100 model instances) while maintaining the magnitude of the individual weights (Table 2, 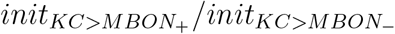). For each model instance, a random number of connections was drawn froma uniform distribution and applied to each MBON (MBON^+^, MBON-) to select the same amount of random connections to be active. While FOC performance remained very robust across all models 1-5 (Figure 5 A), SOC performance was significantly impaired in model 1 and model 3 (Figure 5 B), compared to SOC performance with full connectivity (Figure 3 A).

Additionally, we evaluated the robustness of the learning performance when the strength of the KC>MBON weights was varied in a range of *±*5% around their default weight (Table 2). Again, no significant differences were observed for FOC performance (Figure 5 C). SOC performance was retained for models 2, 4, and 5 compared to the standard model with the same strength of all synaptic weights (Figure 3 A).

While all five circuit motifs are capable of FOC (Figure 2), model 1 performs very poorly at SOC compared to the other models (Figure 2) and also shows poor robustness (Figure 4). While model circuits 3-5 fulfill the criteria for SOC, they differ in their robustness to reward generalization and variations of their parameters. Model 2, which relies on feed-forward input of KCs to the DAN, emerges as a promising candidate, in addition to the circuits that are already being explored [3, 4].

**Figure 4.**
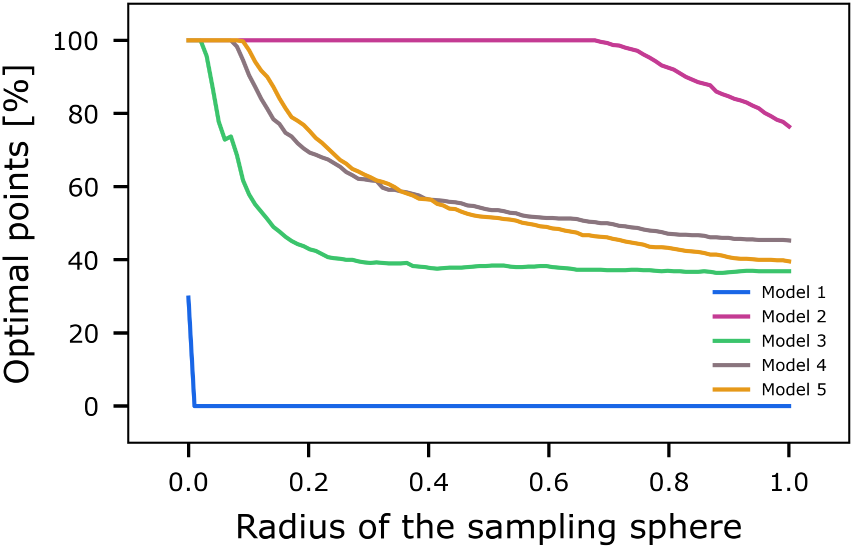
Comparison of parameter robustness between models. The five models were tested for the stability of their second-order learning performance (Eqn. 5) when their optimal parameter combinations were collectively shifted away from their optimum, which we used as the central starting point for a hypersphere with radius=0. We incrementally increased the radius of the hypersphere from 0 to 1 with 100 steps in a linear fashion and sampled 700 data points from its surface for each resulting radius.

**Figure 5.**
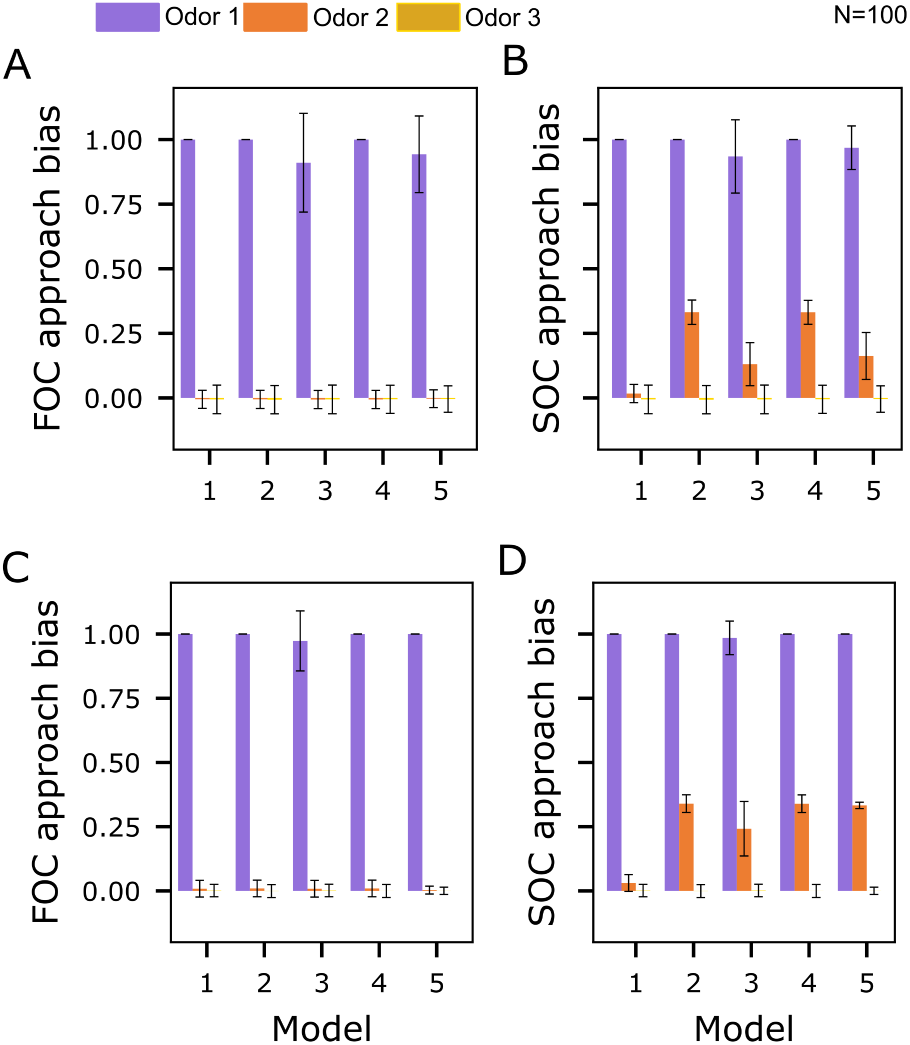
Robustness of the learning performance across variations of the KC>MBON connectivity. (A) Mean approach biases (error bars denote standard deviations, Eqn. 5) to odors 1, 2, and 3 after first order conditioning (FOC) for *N* = 100 model instances with a varying number of KC>MBON synapses, between 25% and 100%, drawn randomly from a uniform distribution. (B) Mean approach biases (Eqn. 5) to odors 1, 2, and 3 after second-order conditioning (SOC) for *N* = 100 model instances with a randomly chosen number of KC>MBON synapses, between 25% and 100%. (C) Mean approach biases (Eqn. 5) to odors 1, 2, and 3 after first order conditioning (FOC) for N=100 model instances with a randomly chosen initial KC>MBON weights (full connectivity), varying *±*5% around the default value (Table 2). (D) Mean approach biases (Eqn. 5) to odors 1, 2, and 3 after second-order conditioning (SOC) for *N* = 100 model instances with a randomly chosen initial KC>MBON weights (full connectivity), varying *±*5% around the default value (Table 2).

## Discussion

While SOC as a phenomenon has been a target of insect learning experiments for a long time [1–3, 5, 6], the discovery of the underlying circuit mechanisms is just at its beginning [2–4]. We use mechanistic models of different variations of a basic, abstract MB circuit inspired by *Drosophila* and show that different circuit motifs, based on either KC or MBON-driven DAN activation, can support SOC. In the following, we will discuss our results in light of experimental evidence for SOC in insects and the extent to which the MB anatomy supports the tested circuit motifs.

### Second-order conditioning in insect experiments and models

Second-order conditioning has been demonstrated experimentally in honeybees [5, 6] and fruit flies [1–4, 56]. The honeybee experiments measured proboscis extension as a response to conditioning with odors and sugar. Regardless of whether the number of SOC trials was equal to [6] or 50% of the number of FOC trials [5], the conditioned response acquired during FOC was stronger than that acquired during SOC. In the fly experiments, a combination of odor and electric shock [1], odor and sugar [3], or odor and optogenetic DAN activation [3] were used. The same duration of pairing an electric shock with odor 1 and pairing odor 1 with odor 2 during SOC yielded a stronger learning effect for FOC, compared to the SOC effect [1], as observed in bees [5, 6]. Yamada et al. [3] used a protocol with longer FOC than SOC duration in an appetitive conditioning protocol. This led to similarly strong FOC and SOC effects, given a long enough FOC training duration.

Potential circuit mechanisms behind SOC were investigated in some studies, conducted in *Drosophila* [2–4]. Evidence for a mechanism based on MBON>DAN feedback comes from a study that used optogenetic silencing or activation of MBONs as an indirect punishment or reward signal for conditioning avoidance or approach of an odor, thereby circumpassing pairing of reinforcement and stimulus 1 during FOC [2]. Yamada et al. [3] also suggest an MBON>DAN pathway across two layers of interneurons as a mechanism for SOC. They show that a presentation of an odor with optogenetic DAN activation can induce suppression of the response of a particular MBON (*α*1). Decreased activity of MBON-*α*1 could cause disinhibition of multiple pathways via two interneurons, leading to a net activation of DANs that encode reward during SOC. The circuit for the disinhibition of the DAN during SOC is closely related to the motif implemented in our model 4, which performs well at SOC but appears not to be very robust to reward generalization onto novel odors. Likewise, in an aversive conditioning paradigm, it was demonstrated that the output of a particular MBON, innervating the *γ*-lobe (MBON-*γ*1), is required during the SOC phase of the experiment to induce a learned valence of the second odor [4]. Similarly, a single MBON (MBON-*α*’2) innervating the *α*’/*β*’-lobes seems to play a similar role in these lobes [4]. In summary, all of these works demonstrate the importance of MBON output pathways [2–4] and seem to suggest MBON>DAN feedback as a candidate mechanism for SOC. For the sake of completeness, it has to be stated that each experiment targeted specific pathways and can not rule out the presence of different circuit motifs for SOC in other compartments.

Recent models of the adult [57, 58] or larval [47] *Drosophila* MB can accommodate SOC. They all suggest KC>MBON plasticity to learn stimulus 2 during SOC via MBON activity. Either in the form of net excitatory and inhibitory MBON>DAN feedback [47, 58] or via direct modulation of the KC>MBON synapses by altered MBON activity [57]. None of the models allow KC input to the DAN and thus do not test this alternative pathway.

### Anatomical evidence for the tested circuit motifs

To evaluate the biological plausibility of our tested circuit motifs, we next assessed which DAN-activation pathways are supported by anatomical evidence. KC>DAN synapses have been found both in larval [46–48] and adult [49–51] *Drosophila*. Since KCs are known to be cholinergic [32] in the adult, it has been suggested that these connections could be excitatory [46] in the larva. This has also been confirmed in adults to affect a particular MBON (*α*2*α*^t^2) [49].

Direct and indirect connections between MBONs and DANs exist in the larva [46, 47] and the adult [51, 52] within and between compartments [46, 47, 51, 52]. In the larva, excitatory and inhibitory synapses have been observed [46, 47]. In the adult *Drosophila* MB, different MBONs have been found to release excitatory or inhibitory transmitters [30], supporting the assumption that here both de- and hyperpolarizing effects of MBONs on DANs might exist.

Direct KC>KC synapses have been found in large numbers in the larva [46]. According to the transmitter released by KCs in the adult [32], they have been speculated to be excitatory [46]. In the adult, KC>KC synapses have also been demonstrated [51, 59]. KCs have been shown to express both muscarinic [59, 60] and nicotinic [61, 62] receptors, the combination of which likely enables both inhibitory [59, 60] and excitatory [63] synapses between them and rendering different interactions plausible.

### Limitations

Motivated by isolating SOC as the phenomenon of interest in our experiments, we decided to reduce our circuit implementations of computational motifs to their minimum and optimize their parameters only for FOC and SOC. This allows us to determine which circuit motifs are the most efficient in computing SOC when optimized solely for this purpose. This approach neglects the surrounding network structures in the real insect MB and thus intentionally disregards other learning phenomena often addressed when studying the MB, such as prediction error, effects of stimulus exposure before learning, forgetting, or extinction. Both forgetting and extinction produce the same observable behavior in experiments, which is a decline in the response to repeated presentation of a sensory cue when reinforcement is omitted after conditioning. As opposed to forgetting, extinction is characterized by the possibility of recovery of the association after its temporary loss [64], has been demonstrated in adult *Drosophila* [65, 66]. To retain the association for recovery, extinction requires the formation of parallel memory traces for the acquisition and the decline of the association [67, 68]. The repeated presentation of stimulus 1 without reinforcement during SOC should lead to the extinction of the learned association between stimulus 1 and reinforcement. Across many trials, SOC and extinction learning should be competing phenomena, allowing SOC to occur only until the extinction process has abolished the odor approach bias. While some studies find a decline in the association between stimulus 1 and the reinforcement during SOC in honeybees [5, 6], experiments in *Drosophila* report no loss of the association between stimulus 1 and reinforcement during SOC [1, 3]. For our *Drosophila*-inspired modeling approach, we thus defined no loss of odor 1 approach bias between the tests following FOC and SOC as a criterion for our parameter optimization. By optimizing the models according to this criterion, we exclude the possibility of studying the effects of loss of an association, such as extinction or forgetting.

Additionally, the omission of mechanisms for long-term network stability in combination with the criterion of perfect odor 1 approach bias in both tests after FOC and SOC guarantee the complete down-regulation of all KC>MBON-synapses activated by odor 1 after three FOC trials of arbitrary duration would not allow for experiments with more trials. Stabilizing mechanisms for homeostasis could be introduced in a more comprehensive modeling approach, implementing the most promising circuit motif in a more naturalistic MB model.

Aside from the narrow applicability to different learning phenomena, which is the downside of our minimal circuit design, another limitation originates from the need to define the limits and the step size for the model parameter grid search. The success of it depends on selecting these limits and steps appropriately (Table 3). If ill-chosen, they could put individual models at a competitive disadvantage by not including or over-stepping their optimum.

### Outlook

We demonstrated that several circuit mechanisms are potential candidates for SOC. While they vary in computational efficiency and robustness, multiple models remain good candidates, compatible with our knowledge of the MB anatomy. A pathway for KC-driven DAN activation emerges as an especially promising candidate among the circuits we tested. A valuable next step would be to integrate them into more comprehensive MB models to test how they interact with other phenomena in learning. This could also be another angle to studying their robustness. Especially interesting in this regard would be extinction, with its inherently interfering mechanism. SOC relies on maintaining the stimulus 1 valence acquired during FOC throughout SOC, which drives DAN activity. Yet, the absence of reinforcement during SOC would trigger the extinction of this very valence. It seems possible that more than one of the circuit motifs could co-exist in different MB compartments. Ultimately, not all MB compartments might be involved in SOC [3, 58], but fulfill other roles in learning.

Computational models are a highly beneficial tool for investigating the circuitry underlying SOC. Experimental validations of theoretically proposed circuit motifs would close the loop between theoretical predictions and their experimental test. However, with the available genetic tools, it is currently impossible to solely manipulate KC>DAN or MBON>DAN synapses either on the pre or post-synaptic side without affecting output onto other or input from other neurons in the network. Therefore, an experimental test of our theoretical predictions is currently difficult to achieve, underlining the importance of computational modeling.

## Methods

### Network Input

Odor and reward signals are provided to all models via the KCs and the DAN. Three odors are used in the experiments. In the most simple case, each exclusively activates an independent combination of (10%) of the 2000 KCs with a rate of 3 Hz to match the levels of population sparseness and low odor-response rates reported in KCs [20, 69, 70]. The first experiment combines the three odors (Figure 1 D). For each model instance, odor 1 activates a combination of 200 randomly chosen KC. KCs activated by odor 2 and odor 3 are then sequentially drawn from the combination of remaining KCs. When odors 1 and 2 are presented together during the experiments, each component of this compound odor activates a random 50% of the KCs, activated by each of the individual odors. This ensures the activated KC does not exceed 10% [19]. The second experiment aims to quantify the stability of the different circuit mechanisms against the generalization of the learned valence onto novel odors. Therefore, a varying degree of odor similarity of odor 3 with odor 1 or odor 2 is used. Odor similarity is implemented as an overlap in activated KCs for odor 1 and odor 2. If any given odor activates 200 KCs, an odor similarity of 50% would yield odors 1 and 2 to have an overlap of 100 KCs. During the joint presentation of both odors, 150 KCs would be activated.

### Parameter Optimization

To increase fairness in the model comparison, all parameters that were part of the grid search were, if possible, optimized within the same boundaries and with the same step size (Table 3). Some parameters were fixed to the same value for all models to adhere to biological constraints (Table 2). KCs show very little spontaneous activity [20, 69, 70] and sparse activation [20, 69–71].

MBON rates of up to 40 Hz have been shown for one MBON [34]. We chose the initial weights for all KC>MBON synapses to achieve plausible MBON rates of no higher than 50 Hz. The upper limit of the DAN rate was 20 Hz to match the experimental literature [55].

A grid search was conducted for each model to optimize the free model parameters (Table 3). All searches contained 100^4^ = 100 10^6^ parameter combinations for testing (Table 3). The grid searches for all models were run on the same server (X86_64 architecture, Ubuntu 20.04.3). The simulation of the parameter combinations was distributed across 24 independent processes using the same random seed. The resulting data were first filtered for the fulfillment of the rate criteria for MBONs and the DAN and the performance in the FOC and SOC tests to determine all *optimal learners*, which meant identifying the parameter combinations with the same coordinates in the objective space (FOC odor 1=1, SOC odor 1=1, SOC odor 2=0.33, and SOC odor 2 0.025 for model 1).

### Hypersphere sampling

Model robustness around the most central optimal parameter combination (see Results: Identifying the optimal parameters for each model) was assessed by testing random samples with increasing distance to the optimal point. We sampled the points **x** uniformly from the surface of a 4-dimensional hypersphere [72, Ch. 1.2.6] with radius *r* by drawing all four components independently of Gaussian distributions with a standard deviation *σ* and scaling with the norm *l***x***l*

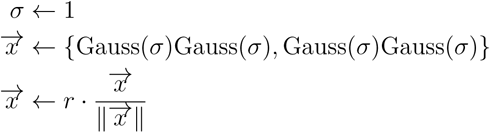

Each sample was evaluated by the indicator function

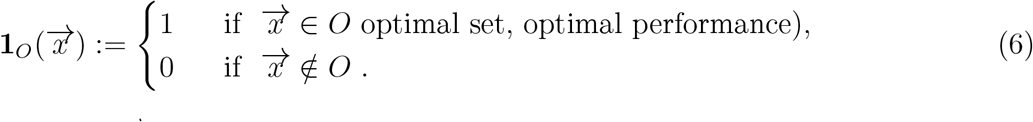

A parameter combination 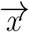 is an element of the optimal set *O* if it shows the same performance in the FOC and SOC tests as the central optimal parameter combination identified in the grid search for the specific model.

The code for implementing the circuit models can be obtained at https://github.com/nawrotlab/exploring_SOC_circuits.

## Acknowledgements

This project is funded in parts by the German Research Foundation (DFG) within the Research Unit ‘Structure, Plasticity and Behavioral Function of the Drosophila mushroom body’ (DFG-FOR-2705, grant no. 403329959, https://www.unigoettingen.de/en/601524.html to MN) and by the Federal Ministry of Education and Research (BMBF, grant no. 01GQ2103A, ‘DrosoExpect’ to MN). AMJ and FS received additional support from the Research Training Group ‘Neural Circuit Analysis’ (DFG-RTG 1960, grant no.233886668). We furthermore thank the Regional Computing Center of the University of Cologne (RRZK) for providing computing time on the DFG-funded (Funding number: INST 216/512/1FUGG) High-Performance Computing (HPC) system CHEOPS. The authors would like to thank André Fiala and El Yazid Rachad for helpful discussions at various project stages and André Fiala for detailed and valuable feedback on the manuscript.

## References

[1] Christopher J Tabone and cond-order conditioning in DrosJ Steven de Belle. “Second-order conditioning in Drosophila”. In: Learning & Memory 18.4 (2011), pp. 250–253. doi: 10.1101/lm.2035411.

[2] Christian König, Afshin Khalili, Thomas Niewalda, Shiqiang Gao, and Bertram Gerber. “An optogenetic analogue of second-order reinforcement in Drosophila”. In: Biology Letters 15.7 (2019), p. 20190084. doi: 10.1098/rsbl.2019.0084.

[3] Daichi Yamada, Daniel Bushey, Feng Li, Karen Hibbard, Megan Sammons, Jan Funke, Ashok Litwin-Kumar, Toshihide Hige, and Yoshinori Aso. “Hierarchical architecture of dopaminergic circuits enables second-order conditioning in Drosophila”. In: bioRxiv (2022). doi: 10.1101/2022.03.30.486484.

[4] El Yazid Rachad. “Neural circuit plasticity underlying learning and memory in Drosophila melanogaster: from synaptic connections to behavior”. PhD thesis. Universität zu Gö”ttingen, 2923. doi: 10.53846/goediss-9845.

[5] ME Bitterman, Randolf Menzel, Andrea Fietz, and Sabine Schäfer. “Classical conditioning of proboscis extension in honeybees (Apis mellifera).” In: Journal of comparative psychology 97.2 (1983), p. 107. doi: 10.1037/0735-7036.97.2.107.

[6] Syed Abid Hussaini, Bernhard Komischke, Randolf Menzel, and Harald Lachnit. “Forward and backward second-order Pavlovian conditioning in honeybees”. In: Learning & Memory 14.10 (2007), pp. 678–683. doi: 10.1101/lm.471307.

[7] Makoto Mizunami, Sae Unoki, Yasuhiro Mori, Daisuke Hirashima, Ai Hatano, and Yukihisa Matsumoto. “Roles of octopaminergic and dopaminergic neurons in appetitive and aversive memory recall in an insect”. In: BMC biology 7.1 (2009), pp. 1–16. doi: 10.1186/1741-7007-7-46.

[8] Robert D Hawkins, Winifred Greene, and Eric R Kandel. “Classical conditioning, differential conditioning, and second-order conditioning of the Aplysia gill-withdrawal reflex in a simplified mantle organ preparation.” In: Behavioral neuroscience 112.3 (1998), p. 636. doi: 10.1037/0735-7044.112.3.636.

[9] Peter C Holland and Robert A Rescorla. “Second-order conditioning with food unconditioned stimulus.” In: Journal of comparative and physiological psychology 88.1 (1975), p. 459. doi: 10.1037/h0076219.

[10] JV Murphy and RE Miller. “Higher-order conditioning in the monkey”. In: The Journal of General Psychology 56.1 (1957), pp. 67–72. doi: 10.1080/00221309.1957.9918363.

[11] Michael Cook and Susan Mineka. “Second-order conditioning and overshadowing in the observational conditioning of fear in monkeys”. In: Behaviour Research and Therapy 25.5 (1987), pp. 349–364. doi: 10.1016/0005-7967(87)90013-1.

[12] Martin Heisenberg. “Mushroom body memoir: from maps to models”. In: Nature Reviews Neuroscience 4.4 (2003), pp. 266–275. doi: 10.1038/nrn1074.

[13] Martin Heisenberg. “What do the mushroom bodies do for the insect brain? An introduction”. In: Learning & memory 5.1 (1998), pp. 1–10. doi: 10.1101/lm.5.1.1.

[14] J Steven De Belle and Martin Heisenberg. “Associative odor learning in Drosophila abolished by chemical ablation of mushroom bodies”. In: Science 263.5147 (1994), pp. 692–695. doi: 10.1126/science.8303280.

[15] Troy Zars. “Behavioral functions of the insect mushroom bodies”. In: Current opinion in neurobiology 10.6 (2000), pp. 790–795. doi: 10.1016/S0959-4388(00)00147-1.

[16] Randolf Menzel and Martin Giurfa. “Cognitive architecture of a mini-brain: the honeybee”. In: Trends in cognitive sciences 5.2 (2001), pp. 62–71. doi: 10.1016/S1364-6613(00)01601-6.

[17] Makoto Mizunami, Josette M Weibrecht, and Nicholas J Strausfeld. “Mushroom bodies of the cockroach: their participation in place memory”. In: Journal of Comparative Neurology 402.4 (1998), pp. 520–537. doi: 10.1002/(SICI)1096-9861(19981228)402:4<520::AID-CNE6>3.0.CO;2-K.

[18] Wolfgang Rössler. “Multisensory navigation and neuronal plasticity in desert ants”. In: Trends in Neurosciences 46.6 (2023), pp. 415–417. doi: 10.1016/j.tins.2023.03.008.

[19] Kyle S Honegger, Robert AA Campbell, and Glenn C Turner. “Cellular-resolution population imaging reveals robust sparse coding in the Drosophila mushroom body”. In: Journal of neuroscience 31.33 (2011), pp. 11772–11785. doi: 10.1523/JNEUROSCI.1099-11.2011.

[20] Glenn C. Turner, Maxim Bazhenov, and Gilles Laurent. “Olfactory representations by Drosophila mushroom body neurons”. In: Journal of Neurophysiology 99.2 (Feb. 2008), pp. 734–746. doi: 10.1152/jn.01283.2007.

[21] Andrew C Lin, Alexei M Bygrave, Alix De Calignon, Tzumin Lee, and Gero Miesenböck. “Sparse, decorrelated odor coding in the mushroom body enhances learned odor discrimination”. In: Nature neuroscience 17.4 (2014), pp. 559–568. doi: 10.1038/nn.3660.

[22] Makoto Mizunami, Masayuki Iwasaki, Michiko Nishikawa, and Ryuichi Okada. “Modular structures in the mushroom body of the cockroach”. In: Neuroscience letters 229.3 (1997), pp. 153–156. doi: 10.1016/S0304-3940(97)00438-2.

[23] Makoto Mizunami, Masayuki Iwasaki, Ryuichi Okada, and Michiko Nishikawa. “Topography of modular subunits in the mushroom bodies of the cockroach”. In: Journal of Comparative Neurology 399.2 (1998), pp. 153–161. doi: 10.1002/(SICI)1096-9861(19980921)399:2<153::AID-CNE1>3.0.CO;2-%23.

[24] Jürgen Rybak and Randolf Menzel. “Anatomy of the mushroom bodies in the honey bee brain: the neuronal connections of the alpha-lobe”. In: Journal of Comparative Neurology 334.3 (1993), pp. 444–465. doi: 10.1002/cne.903340309.

[25] Susan E Fahrbach. “Structure of the mushroom bodies of the insect brain”. In: Annu. Rev. Entomol. 51 (2006), pp. 209–232. doi: 10.1146/annurev.ento.51.110104.150954.

[26] Yoshinori Aso, Daisuke Hattori, Yang Yu, Rebecca M Johnston, Nirmala A Iyer, Teri-TB Ngo, Heather Dionne, LF Abbott, Richard Axel, Hiromu Tanimoto, et al. “The neuronal architecture of the mushroom body provides a logic for associative learning”. In: elife 3 (2014), e04577. doi: 10.7554/eLife.04577.

[27] Nobuaki K Tanaka, Hiromu Tanimoto, and Kei Ito. “Neuronal assemblies of the Drosophila mushroom body”. In: Journal of Comparative Neurology 508.5 (2008), pp. 711–755. doi: 10.1002/cne.21692.

[28] Martin F Strube-Bloss and Wolfgang Rössler. “Multimodal integration and stimulus categorization in putative mushroom body output neurons of the honeybee”. In: Royal Society Open Science 5.2 (2018), p. 171785. doi: 10.1098/rsos.171785.

[29] Yongsheng Li and Nicholas J Strausfeld. “Morphology and sensory modality of mushroom body extrinsic neurons in the brain of the cockroach, Periplaneta americana”. In: Journal of Comparative Neurology 387.4 (1997), pp. 631–650. doi: 10.1002/(SICI)1096-9861(19971103)387:4<631::AID-CNE9>3.0.CO;2-3.

[30] Yoshinori Aso, Divya Sitaraman, Toshiharu Ichinose, Karla R Kaun, Katrin Vogt, Ghislain Belliart-Guérin, Pierre-Yves Plaçais, Alice A Robie, Nobuhiro Yamagata, Christopher Schnaitmann, et al. “Mushroom body output neurons encode valence and guide memory-based action selection in Drosophila”. In: Elife 3 (2014), e04580. doi: /10.7554/eLife.04580.

[31] David Owald and Scott Waddell. “Olfactory learning skews mushroom body output pathways to steer behavioral choice in Drosophila”. In: Current opinion in neurobiology 35 (2015), pp. 178–184. doi: /10.1016/j.conb.2015.10.002.

[32] Oliver Barnstedt, David Owald, Johannes Felsenberg, Ruth Brain, John-Paul Moszynski, Clifford B Talbot, Paola N Perrat, and Scott Waddell. “Memory-relevant mushroom body output synapses are cholinergic”. In: Neuron 89.6 (2016), pp. 1237–1247. doi: 10.1016/j.neuron.2016.02.015.

[33] David Owald, Johannes Felsenberg, Clifford B Talbot, Gaurav Das, Emmanuel Perisse, Wolf Huetteroth, and Scott Waddell. “Activity of defined mushroom body output neurons underlies learned olfactory behavior in Drosophila”. In: Neuron 86.2 (2015), pp. 417–427. doi: 10.1016/j.neuron.2015.03.025.

[34] Toshihide Hige, Yoshinori Aso, Mehrab N Modi, Gerald M Rubin, and Glenn C Turner. “Heterosynaptic plasticity underlies aversive olfactory learning in Drosophila”. In: Neuron 88.5 (2015), pp. 985–998. doi: /10.1016/j.neuron.2015.11.003.

[35] Martin F Strube-Bloss, Martin P Nawrot, and Randolf Menzel. “Mushroom body output neurons encode odor–reward associations”. In: Journal of Neuroscience 31.8 (2011), pp. 3129–3140. doi: 10.1523/JNEUROSCI.2583-10.2011.

[36] Yoshinori Aso and Gerald M Rubin. “Dopaminergic neurons write and update memories with cell-type-specific rules”. In: Elife 5 (2016), e16135. doi: /10.7554/eLife.16135.

[37] Scott Waddell. “Dopamine reveals neural circuit mechanisms of fly memory”. In: Trends in neurosciences 33.10 (2010), pp. 457–464. doi: /10.1016/j.tins.2010.07.001.

[38] Silke Sachse and Jennifer Beshel. “The good, the bad, and the hungry: how the central brain codes odor valence to facilitate food approach in Drosophila”. In: Current opinion in neurobiology 40 (2016), pp. 53–58. doi: 10.1016/j.conb.2016.06.012.

[39] Scott Waddell. “Reinforcement signalling in Drosophila; dopamine does it all after all”. In: Current opinion in neurobiology 23.3 (2013), pp. 324–329. doi: 10.1016/j.conb.2013.01.005.

[40] Young-Cho Kim, Hyun-Gwan Lee, Junghwa Lim, and Kyung-An Han. “Appetitive learning requires the alpha1-like octopamine receptor OAMB in the Drosophila mushroom body neurons”. In: Journal of Neuroscience 33.4 (2013), pp. 1672–1677. doi: 10.1523/JNEUROSCI.3042-12.2013.

[41] Martin Schwaerzel, Maria Monastirioti, Henrike Scholz, Florence Friggi-Grelin, Serge Birman, and Martin Heisenberg. “Dopamine and octopamine differentiate between aversive and appetitive olfactory memories in Drosophila”. In: Journal of Neuroscience 23.33 (2003), pp. 10495–10502. doi: 10.1523/JNEUROSCI.23-33-10495.2003.

[42] Makoto Mizunami and Yukihisa Matsumoto. “Roles of octopamine and dopamine neurons for mediating appetitive and aversive signals in Pavlovian conditioning in crickets”. In: Frontiers in physiology 8 (2017), p. 1027. doi: 10.3389/fphys.2017.01027.

[43] Kristina V Dylla, Georg Raiser, C Giovanni Galizia, and Paul Szyszka. “Trace conditioning in Drosophila induces associative plasticity in mushroom body Kenyon cells and dopaminergic neurons”. In: Frontiers in neural circuits 11 (2017), p. 42. doi: 10.3389/fncir.2017.00042.

[44] Thomas Riemensperger, Thomas Völler, Patrick Stock, Erich Buchner, and André Fiala. “Punishment prediction by dopaminergic neurons in Drosophila”. In: Current Biology 15.21 (2005), pp. 1953–1960. doi: 10.1016/j.cub.2005.09.042.

[45] Wolfram Schultz. “Predictive reward signal of dopamine neurons”. In: Journal of neurophysiology (1998). doi: 10.1152/jn.1998.80.1.1.

[46] Katharina Eichler, Feng Li, Ashok Litwin-Kumar, Youngser Park, Ingrid Andrade, Casey M Schneider-Mizell, Timo Saumweber, Annina Huser, Claire Eschbach, Bertram Gerber, Richard D Fetter, James W Truman, Carey E Priebe, L F Abbott, Andreas S Thum, Marta Zlatic, and Albert Cardona. “The complete connectome of a learning and memory centre in an insect brain”. In: Nature 548.7666 (2017), pp. 175–182. doi: 10.1038/nature23455.

[47] Claire Eschbach, Akira Fushiki, Michael Winding, Casey M Schneider-Mizell, Mei Shao, Rebecca Arruda, Katharina Eichler, Javier Valdes-Aleman, Tomoko Ohyama, Andreas S Thum, Bertram Gerber, Richard D Fetter, James W Truman, Ashok Litwin-Kumar, Albert Cardona, and Marta Zlatic. “Recurrent architecture for adaptive regulation of learning in the insect brain”. In: Nature Neuroscience 23.4 (2020), pp. 544–555. doi: 10.1038/s41593-020-0607-9.

[48] Timo Saumweber, Astrid Rohwedder, Michael Schleyer, Katharina Eichler, Yi-Chun Chen, Yoshinori Aso, Albert Cardona, Claire Eschbach, Oliver Kobler, Anne Voigt, Archana Durairaja, Nino Mancini, Marta Zlatic, James W Truman, Andreas S Thum, and Bertram Gerber. “Functional architecture of reward learning in mushroom body extrinsic neurons of larval Drosophila”. In: Nature communications 9.1 (2018), p. 1104. doi: 10.1038/s41467-018-03130-1.

[49] Isaac Cervantes-Sandoval, Anna Phan, Molee Chakraborty, and Ronald L Davis. “Reciprocal synapses between mushroom body and dopamine neurons form a positive feedback loop required for learning”. In: Elife 6 (2017), e23789. doi: /10.7554/eLife.23789.

[50] Louis K Scheffer, C Shan Xu, Michal Januszewski, Zhiyuan Lu, Shin-ya Takemura, Kenneth J Hayworth, Gary B Huang, Kazunori Shinomiya, Jeremy Maitlin-Shepard, Stuart Berg, et al. “A connectome and analysis of the adult Drosophila central brain”. In: Elife 9 (2020), e57443. doi:/10.7554/eLife.57443.

[51] Shin-ya Takemura, Yoshinori Aso, Toshihide Hige, Allan Wong, Zhiyuan Lu, C Shan Xu, Patricia K Rivlin, Harald Hess, Ting Zhao, Toufiq Parag, et al. “A connectome of a learning and memory center in the adult Drosophila brain”. In: Elife 6 (2017), e26975. doi: 10.7554/eLife.26975.

[52] Toshiharu Ichinose, Yoshinori Aso, Nobuhiro Yamagata, Ayako Abe, Gerald M Rubin, and Hiromu Tanimoto. “Reward signal in a recurrent circuit drives appetitive long-term memory formation”. In: Elife 4 (2015), e10719.

[53] Julien Séjourné, Pierre-Yves Plaçais, Yoshinori Aso, Igor Siwanowicz, Séverine Trannoy, Vladimiros Thoma, Stevanus R Tedjakumala, Gerald M Rubin, Paul Tchénio, Kei Ito, et al. “Mushroom body efferent neurons responsible for aversive olfactory memory retrieval in Drosophila”. In: Nature neuroscience 14.7 (2011), pp. 903–910. doi: /10.1038/nn.2846.

[54] Clare E Hancock, Vahid Rostami, El Yazid Rachad, Stephan H Deimel, Martin P Nawrot, and André Fiala. “Visualization of learning-induced synaptic plasticity in output neurons of the Drosophila mushroom body γ-lobe”. In: Scientific Reports 12.1 (2022), p. 10421. doi: 10.1038/s41598-022-14413-5.

[55] Cheng Huang, Junjie Luo, Seung Je Woo, Lucas Roitman, Jizhou Li, Vincent Pieribone, Madhuvanthi Kannan, Ganesh Vasan, and Mark Schnitzer. “Dopamine signals integrate innate and learnt valences to regulate memory dynamics”. In: Research Square (2022). doi: 10.21203/rs.3.rs-1915648/v1.

[56] Björn Brembs and Martin Heisenberg. “Conditioning with compound stimuli in Drosophila melanogaster in the flight simulator”. In: Journal of experimental biology 204.16 (2001), pp. 2849–2859. doi: 10.1242/jeb.204.16.2849.

[57] Faramarz Faghihi, Ahmed A Moustafa, Ralf Heinrich, and Florentin Wörgötter. “A computational model of conditioning inspired by Drosophila olfactory system”. In: Neural Networks 87 (2017), pp. 96–108. doi: 10.1016/j.neunet.2016.11.002.

[58] Linnie Jiang and Ashok Litwin-Kumar. “Models of heterogeneous dopamine signaling in an insect learning and memory center”. In: PLoS Computational Biology 17.8 (2021), e1009205. doi: /10.1371/journal.pcbi.1009205.

[59] Julia E Manoim, Andrew M Davidson, Shirley Weiss, Toshihide Hige, and Moshe Parnas. “Lateral axonal modulation is required for stimulus-specific olfactory conditioning in Drosophila”. In: Current Biology 32.20 (2022), pp. 4438–4450. doi: 10.1016/j.cub.2022.09.007.

[60] Noa Bielopolski, Hoger Amin, Anthi A Apostolopoulou, Eyal Rozenfeld, Hadas Lerner, Wolf Huetteroth, Andrew C Lin, and Moshe Parnas. “Inhibitory muscarinic acetylcholine receptors enhance aversive olfactory learning in adult Drosophila”. In: eLife 8 (2019), e48264. doi: 10.7554/eLife.48264.

[61] Amanda Crocker, Xiao-Juan Guan, Coleen T. Murphy, and Mala Murthy. “Cell-Type-Specific Transcriptome Analysis in the Drosophila Mushroom Body Reveals Memory-Related Changes in Gene Expression”. In: Cell Reports 15.7 (2016), pp. 1580–1596. doi: 10.1016/j.celrep.2016.04.046.

[62] Vincent Croset, Christoph D Treiber, and Scott Waddell. “Cellular diversity in the Drosophila midbrain revealed by single-cell transcriptomics”. In: eLife 7 (2018), e34550. doi: 10.7554/eLife.34550.

[63] Hailing Su and Diane K O’Dowd. “Fast synaptic currents in Drosophila mushroom body Kenyon cells are mediated by α-bungarotoxin-sensitive nicotinic acetylcholine receptors and picrotoxin-sensitive GABA receptors”. In: (). doi: 10.1523/JNEUROSCI.23-27-09246.2003.

[64] Mark E Bouton. “Context and behavioral processes in extinction”. In: Learning & memory 11.5 (2004), pp. 485–494. doi: 10.1101/lm.78804.

[65] Lingling Wang, Qi Yang, Binyan Lu, Lianzhang Wang, Yi Zhong, and Qian Li. “A behavioral paradigm to study the persistence of reward memory extinction in Drosophila”. In: Journal of genetics and genomics 46.12 (2019), pp. 599–601. doi: 10.1016/j.jgg.2019.11.001.

[66] Yukinori Hirano, Kunio Ihara, Tomoko Masuda, Takuya Yamamoto, Ikuko Iwata, Aya Takahashi, Hiroko Awata, Naosuke Nakamura, Mai Takakura, Yusuke Suzuki, Junjiro Horiuchi, Hiroyuki Okuno, and Minoru Saitoe. “Shifting transcriptional machinery is required for longterm memory maintenance and modification in Drosophila mushroom bodies”. In: Nature communications 7.1 (2016), p. 13471. doi: 10.1038/ncomms13471.

[67] Johannes Felsenberg, Oliver Barnstedt, Paola Cognigni, Suewei Lin, and Scott Waddell. “Reevaluation of learned information in Drosophila”. In: Nature 544.7649 (2017), pp. 240–244. doi: 10.1038/nature21716.

[68] Johannes Felsenberg, Pedro F Jacob, Thomas Walker, Oliver Barnstedt, Amelia J Edmondson-Stait, Markus W Pleijzier, Nils Otto, Philipp Schlegel, Nadiya Sharifi, Emmanuel Perisse, et al. “Integration of parallel opposing memories underlies memory extinction”. In: Cell 175.3 (2018), pp. 709–722. doi: /10.1016/j.cell.2018.08.021.

[69] Javier Perez-Orive, Ofer Mazor, Glenn C Turner, Stijn Cassenaer, Rachel I Wilson, and Gilles Laurent. “Oscillations and sparsening of odor representations in the mushroom body”. In: Science 297.5580 (2002), pp. 359–365. doi: 10.1126/science.1070502.

[70] Iori Ito, Rose Chik-Ying Ong, Baranidharan Raman, and Mark Stopfer. “Sparse odor representation and olfactory learning”. In: Nature neuroscience 11.10 (2008), pp. 1177–1184. doi: 10.1038/nn.2192.

[71] Eleftheria Vrontou, Lukas N Groschner, Susanne Szydlowski, Ruth Brain, Alina Krebbers, and Gero Miesenböck. “Response competition between neurons and antineurons in the mushroom body”. In: Current Biology 31.22 (2021), pp. 4911–4922. doi: 10.1016/j.cub.2021.09.008.

[72] Werner Krauth. Statistical mechanics : algorithms and computations. Oxford master series in statistical, computational, and theoretical physics. Oxford: Oxford University Press, 2006.

